# Identification of a lymphocyte minor histocompatibility antigen in Mauritian cynomolgus macaques

**DOI:** 10.1101/2020.06.10.145250

**Authors:** Jason T. Weinfurter, Michael E. Graham, Adam J. Ericsen, Lea M. Matschke, Sian Llewellyn-Lacey, David A. Price, Roger W. Wiseman, Matthew R. Reynolds

**Affiliations:** Department of Pathobiological Sciences, School of Veterinary Medicine, University of Wisconsin-Madison, Madison, Wisconsin, USA; Wisconsin National Primate Research Center, University of Wisconsin-Madison, Madison, Wisconsin, USA; Division of Infection and Immunity, Cardiff University School of Medicine, Cardiff, Wales, United Kingdom; Systems Immunity Research Institute, Cardiff University School of Medicine, Cardiff, Wales, UK

## Abstract

Allogeneic hematopoietic stem cell transplantation can lead to dramatic reductions in human immunodeficiency virus (HIV) reservoirs. This effect is mediated in part by donor T cells that recognize lymphocyte-expressed minor histocompatibility antigens (mHAgs). The potential to mark malignant and latently infected cells for destruction makes mHAgs attractive targets for cellular immunotherapies. However, testing such HIV reservoir reduction strategies will likely require preclinical studies in nonhuman primates (NHPs). In this study, we used a combination of alloimmunization, whole exome sequencing, and bioinformatics to identify a mHAg in Mauritian cynomolgus macaques (MCMs). We mapped the minimal optimal epitope to a 10-mer peptide (SW10) in apolipoprotein B mRNA editing enzyme catalytic polypeptide-like 3C (APOBEC3) and determined the major histocompatibility complex class I restriction element as Mafa-A1*063, which is expressed in almost 90% of MCMs. APOBEC3C SW10-specific CD8+ T cells recognized immortalized B cells but not fibroblasts from a mHAg positive MCM. These results collectively provide a framework for identifying mHAgs in a nontransplant setting and suggest that APOBEC3C SW10 could be used as a lymphocyte-restricted model antigen in NHPs to test various mHAg-targeted immunotherapies.

**Importance:** Cellular immunotherapies developed to treat blood cancers may also be effective against latent HIV. Preclinical studies of such immunotherapies are hindered by a lack of known target antigens. We used a combination of alloimmunization, basic immune assays, whole exome sequencing, and bioinformatics to identify a lymphocyte-restricted minor histocompatibility antigen in a genetically related population of nonhuman primates. This minor histocompatibility antigen provides an actionable target for piloting cellular immunotherapies designed to reduce or eliminate latent reservoirs of HIV.

## Background

The establishment of long-lived viral reservoirs after primary infection is a major obstacle to the development of curative therapies for human immunodeficiency virus (HIV) (1–4). These latent reservoirs are unaffected by antiretroviral therapy (ART) and readily reactivate upon cessation of treatment (5, 6). Moreover, it has been estimated that latently infected cells, which exist at low frequencies *in vivo*, would only decay to extinction after more than 60 years in the presence of continuous ART (2, 7). Strategies are therefore being developed to reduce or eliminate viral reservoirs and attain sustained ART-free remission (8, 9).

Dramatic reductions in latent viral reservoirs have been observed in HIV positive patients with hematological malignancies undergoing treatment with allogeneic hematopoietic stem cell transplants (allo-HSCTs). In two widely publicized cases, the “Berlin” and “London” patients achieved ART-free remission after receiving allo-HSCTs from donors homozygous for the *CCR5Δ32* gene mutation (10, 11). The absence of functional CCR5 co-receptors on the surface of donor-derived immune cells is undoubtedly a crucial factor in the remarkable success of these interventions, preventing the reseeding of the newly established hematopoietic system. However, graft-versus-host (GvH) responses likely contributed to the elimination of endogenous viral reservoirs (10, 12). Interestingly, the London patient received reduced-intensity conditioning prior to allo-HSCT, leaving a residual pool of cancerous and HIV latently infected cells (10, 13). In this setting, allogeneic T cell responses potentially eliminated both malignant and latently infected cells by simultaneously mediating “graft-versus-leukemia” and “graft-versus-HIV” effects. Intriguingly, similar reductions in latent viral reservoirs have been achieved using HSCs from donors expressing wild-type CCR5 (14–16). The most notable examples are the “Boston” patients, who received allo-HSCTs under cover of ART and exhibited prolonged ART-fee remission before viral rebound (14). Collectively, these anecdotal cases demonstrate that alloreactive T cells can attack and destroy endogenous hematopoietic cells latently infected with HIV.

Alloreactive T cells commonly recognize minor histocompatibility antigens (mHAgs). mHAgs are polymorphic peptides presented by major histocompatibility complex (MHC) molecules and recognized as “foreign” by allogeneic T cells (17). The tissue distribution and expression profile of mHAg-encoding genes affect the outcome of GvH responses (18, 19). Expression of mHAgs across a broad range of tissues can result in toxic GvH reactions and potentially fatal GvH disease (GvHD). Conversely, mHAgs presented exclusively by recipient leukocytes, including malignant and latently infected cells, can elicit beneficial GvH responses that eliminate tumors and latently infected cells alike. Immunotherapies targeting leukocyte-derived mHAgs may therefore provide a novel approach to eliminating hematopoietic cells latently infected with HIV. Further studies are required to advance this concept, however, and the potential efficacy of graft-versus-HIV responses can only realistically be assessed in nonhuman primates (NHPs) (20).

Nonhuman primates (NHP) are common models in preclinical organ transplant and infectious disease studies. Despite their ubiquitous use in research, no mHAgs have been identified in NHPs. In contrast, several approaches have been used to identify various human mHAgs including peptide elution, cDNA library screens, and genetic linkage analyses using large panels of immortalized B cells (21). Recently, these labor-intensive methods have been complemented by advances in next-generation sequencing technologies to accelerate the pace of mHAg discovery (22).

Most outbred populations of NHPs are genetically diverse, making it challenging to identify cohorts of animals with one or more matches at the MHC class I (MHC-I) locus. In contrast, Mauritian-origin cynomolgus macaques (MCM) are descended from a small founder population approximately 400 years ago (23), resulting in limited genetic diversity, even within highly polymorphic loci (24–26). Indeed, only seven major MHC haplotypes, designated M1 through M7, have been identified in MCMs.

The unique population genetics of MCM provides an opportunity to map mHAgs in a relevant NHP transplant and HIV cure model. In this report, we describe the identification and characterization of a mHAg, APOBEC3C SW10, in MCM. This epitope is restricted by the MHC-I molecule Mafa-A1*063, which is expressed by almost 90% of MCMs (27). Our data suggests that the APOBEC3C SW10 may serve as a useful model antigen for interventional studies of mHAg-targeted therapies in NHPs.

## Results

### Generation of mHAg-specific CD8^+^ T cell clones

We identified 21 MCM heterozygous for the M1 and M2 MHC haplotypes (M1/M2). mHAg-reactive T cells were stimulated *in vivo* by alloimmunizing four M1/M2 MCMs (vaccinees) with a mixture of peripheral blood mononuclear cells (PBMC) from four MHC-identical MCM (donors). Allo-reactive T cells were then isolated by incubating PBMCs from the vaccinees with a mixture of irradiated immortalized donor B lymphoblastoid cell lines (BLCLs). After 3 weeks, these T-cell lines reacted to allogeneic but not autologous BLCLs in IFN-γ ELISpot assays, indicating specific recognition of donor-restricted mHAgs.

To identify individual mHAgs, we isolated T cell clones via limiting dilution of the alloreactive T cell lines. After 8 weeks in culture, these T cell clones (n=60) displayed unique patterns of binary reactivity against a panel of MHC-matched allogeneic BLCLs (n=10) in IFN-γ ELISpot assays, indicating specific recognition of distinct mHAgs (Figure 1). These reactivity patterns were used to identify genomic polymorphisms associated with potential mHAgs.

**Figure 1.**
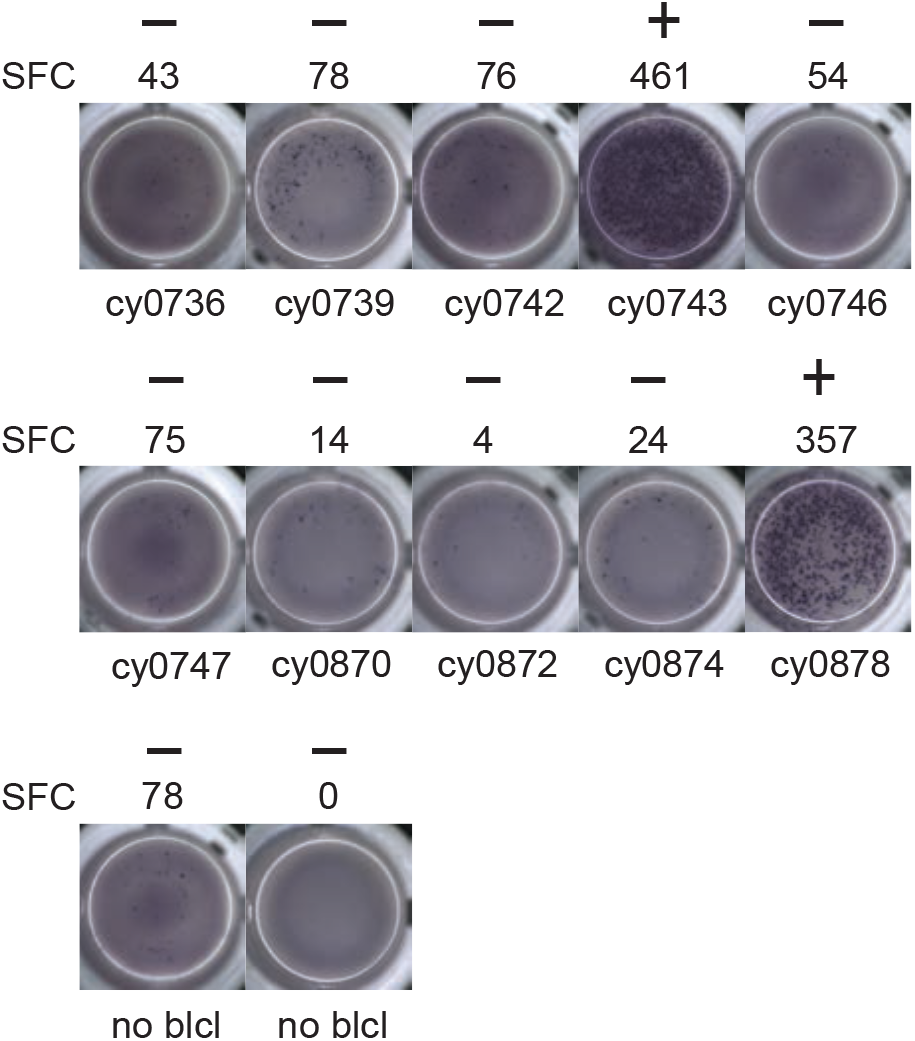
Alloreactive T cell clones differentially recognize allogeneic BLCL. Representative data from T cell clone 11 demonstrating strong reactivity against two of the ten allogeneic BLCLs. Equal numbers of T cells and BLCLs were incubated overnight in IFN-γ ELISpot assay. (+) and (−) signify animals that were placed into mHAg^pos^ and mHAg^neg^ groups for WES segregation analysis, respectively. SFC, spot-forming cell.

### Identification of mHAg-associated SNPs in APOBEC3C

We reasoned that comparing the genomic sequences of MCM whose BLCLs were or were not recognized by alloreactive T cell clones would simplify the winnowing of germline polymorphisms associated with mHAgs. Accordingly, we performed whole exome sequencing (WES) using a probe set specially designed to capture MCM coding sequences, which enabled the generation of single nucleotide polymorphism (SNP) profiles for each MCM. Using a custom-built analysis tool, we then parsed the WES data for non-synonymous SNPs (ns-SNPs) that distinguished T cell clone-specific mHAg^pos^ and mHAg^neg^ groups of BLCLs. We prioritized ns-SNPs present only in the mHAg^pos^ group for further analysis, surmising these ns-SNPs likely encoded the relevant mHAgs.

We focused our mHAg mapping efforts on the ns-SNPs associated with T cell clones 2, 4, 11, and 14. The segregation analysis identified SNPs encoding amino acid polymorphisms in the *IGHM, OR4K3, APOBEC3C*, and *IL20RA/MAP3K5* genes. For each candidate mHAg, we identified the 11 amino acids upstream and the 11 amino acids downstream of the corresponding SNPs and synthesized 15-mer peptides overlapping by 11 amino acids. Each 15-mer peptide was then pulsed onto mHAg^neg^ BLCLs at a concentration of 1 μM and incubated with their respective T cell clone. In IFN-γ ELISpot assays, T cell clone 11 (T11) strongly recognized two of the three 15-mers containing an arginine-to-leucine amino acid change, relative to the mHAg^neg^ sequence, in APOBEC3C (Figure 2A). The other 15-mer peptides failed to elicit IFN-γ production in parallel assays with T2, T4, or T14. We therefore concentrated on identifying the minimal optimal epitope recognized by T11.

**Figure 2.**
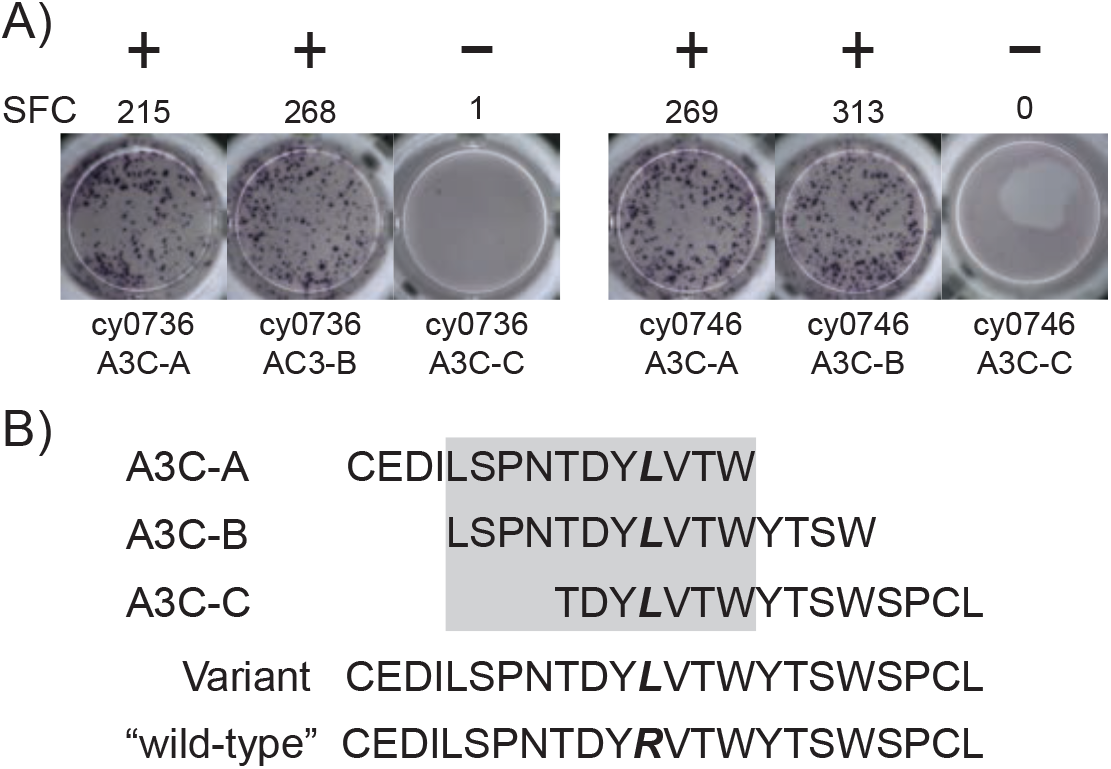
An mHAg-reactive T cell clone recognizes an amino acid variant in APOBEC3C. (A) T cell clone T11 was incubated with mHAg^neg^ BLCLs from cy0736 (left) or cy0746 (right) pulsed with overlapping 15-mer peptides containing an arginine-to-leucine amino acid change relative to the mHAg^neg^ sequence in an IFN-γ ELISpot assay. (B) Sequences of the APOBEC3C (A3C) peptides used to pulse the mHAg^neg^ BLCL in (A). The variant amino acid is highlighted in bold. The grey box corresponds to the putative region containing the mHAg epitope. (+) and (−) signify wells determined to be positive or negative, respectively. SFC, spot-forming cell.

### Identification of the minimal optimal mHAg epitope in APOBEC3C

We reasoned that the mHAg epitope recognized by T11 was present in each of the two stimulatory APOBEC3C 15-mer peptides, denoted as A3C-A and A3C-B, but not in the non-stimulatory APOBEC3C 15-mer peptide, denoted as A3C-C (Figure 2B). Accordingly, we synthesized a series of overlapping 8-mer, 9-mer, 10-mer, and 11-mer peptides spanning the 11 amino acids present in A3C-A and A3C-B to map the minimal optimal epitope in APOBEC3C. We also synthesized a 15-mer and 11-mer peptide matching the reference, “wild-type”, sequence with an arginine replacing the variant leucine (Figure 2B). In intracellular cytokine staining (ICS) assays, a bulk T cell line generated against A3C-A and A3C-B strongly recognized the 11-mer peptide LW11, which matched the intersecting region of the overlapping 15-mer peptides (Figure 3). A similar response was observed with the 10-mer peptide SW10. In contrast, weaker responses were observed with all other peptides, including LT10 and the wild-type peptides, which induced IFN-γ production from approximately a third as many T cells as the corresponding variant peptides.

**Figure 3.**
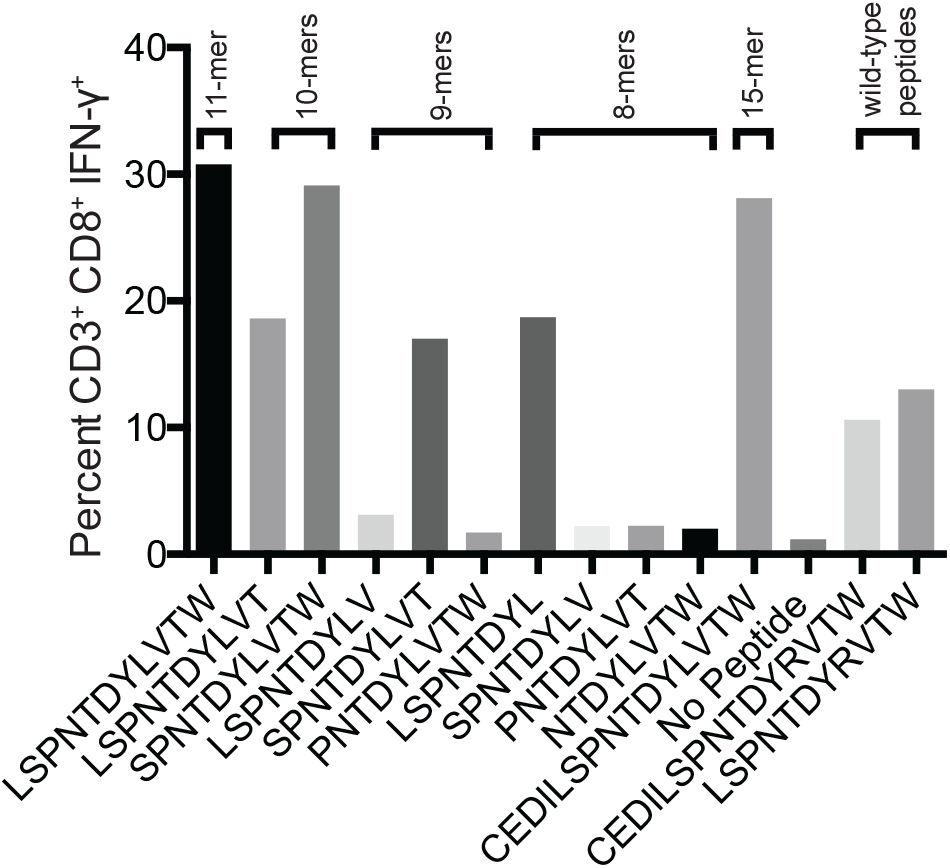
Select peptides corresponding to the APOBEC3C variant stimulate an mHAg-specific T cell line. An APOBEC3C mHAg-specific T cell line was incubated with mHAg^neg^ BLCL pulsed with peptides, including the 11-mer peptide corresponding to the shared region between the 15-mer peptides A3C-A and A3C-B (Figure 2C) and corresponding “wild-type” 15-mer and 11-mer peptides. Unpulsed autologous BLCLs were included as a negative control, and A3C-A peptide-pulsed autologous BLCLs were included as a negative control. Displayed is the percentage of CD3^+^CD8^+^IFN-γ^+^ determined by ICS.

To confirm these results, we conducted similar ICS assays using serial dilutions of each peptide that elicited a response at a concentration of 1 μM, namely LL8, ST9, LT10, SW10, and LW11. Sharp reductions in IFN-γ were observed with decreasing concentrations of peptides LL8, ST9, and LT10 (Figure 4A). In contrast, SW10 and LT11 elicited similar levels of IFN-γ production throughout the dilution series, which extended down to 1 nM. In further titration experiments, SW10 and LT11 exhibited largely equivalent dose-response curves down to a concentration of 1 fM, but SW10 elicited more potently at concentrations of 1 nM and 100 pM, suggesting this peptide was the minimal optimal mHAg epitope in APOBEC3C (Figure 4B).

**Figure 4.**
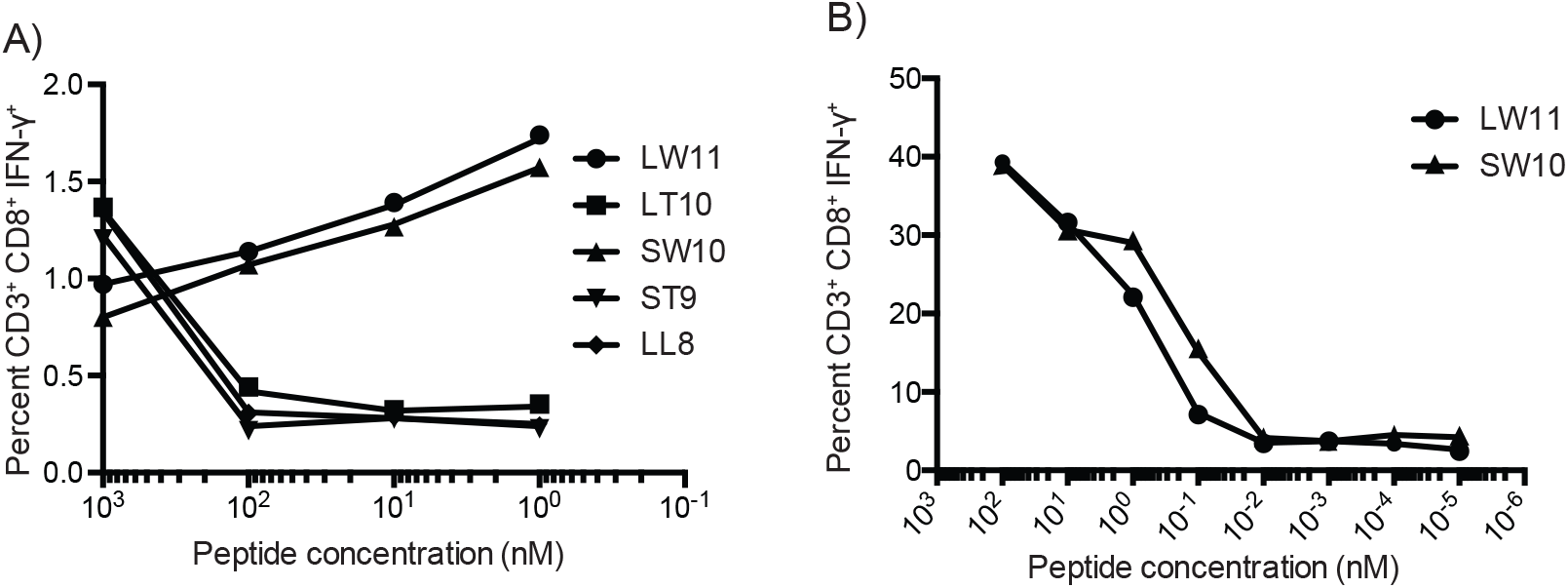
Fine mapping of the mHAg epitope in APOBEC3C. (A) A APOBEC3C mHAg-specific T cell line was incubated with mHAg^neg^ BLCLs pulsed with serial dilutions of the indicated peptides from 1μM to 1nM. (B) As in (A) with serial dilutions of the peptides SW10 and LW11 from 100nM to 1fM. Displayed is the percentage of CD3^+^CD8^+^IFN-γ^+^ determined by ICS.

### MHC class I restriction of the mHAg epitope in APOBEC3C

To determine the restricting MHC-I molecule for APOBEC3C SW10, we used three immortalized human MHC-I null cell lines individually expressing Mafa-A and Mafa-B allomorphs from the MCM M2 MHC haplotype (28). Each of these MHC-I “transferents” was pulsed with the SW10 peptide and incubated with the APOBEC3C-specific T cell line. In ICS assays, only the T cells incubated with the peptide-pulsed Mafa-A1*063 transferent induced IFN-γ production above background levels, indicating stable complexes with SW10 (Figure 5).

**Figure 5.**
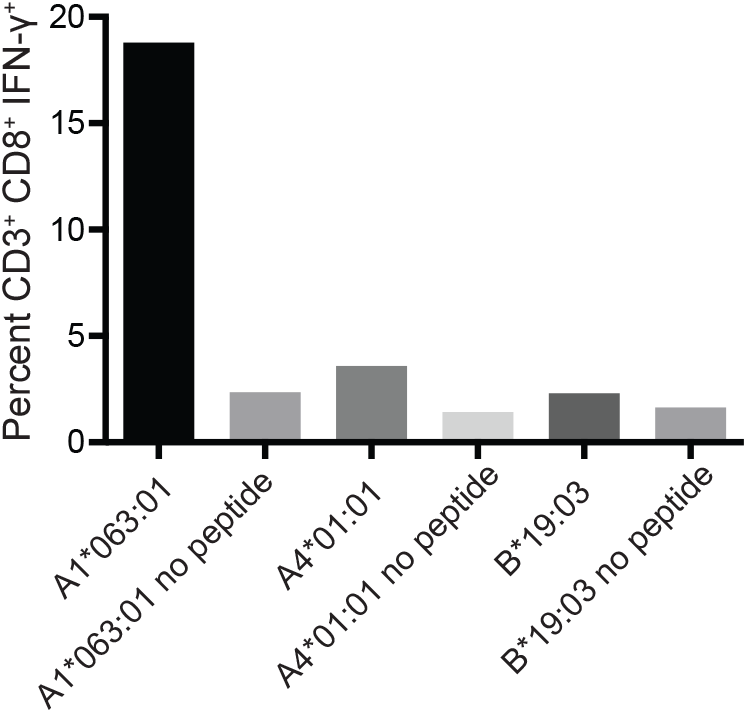
Restriction analysis of the mHAg epitope in APOBEC3C. An APOBEC3C mHAg-specific T cell line was incubated with unpulsed or SW10 peptide-pulsed MHC-I transferents expressing Mafa-A1*063:01, Mafa-A4:01:01, or Mafa-B*19:03. Displayed is the percentage of CD3^+^CD8^+^IFN-γ^+^ determined by ICS.

The SW10 epitope corresponds to the previously described Mafa-A1*063 peptide-binding motif, with a serine at position one, a proline at position two, an asparagine at position three, and a tryptophan at the carboxy terminus of the peptide (24, 29). We were therefore able to form tetrameric complexes of Mafa-A1*063/SW10. In line with the functional data, these recombinant antigen complexes robustly stained APOBEC3C-specific T cells, confirming Mafa-A1*063 as the restricting element for SW10 (Figure 6).

**Figure 6.**
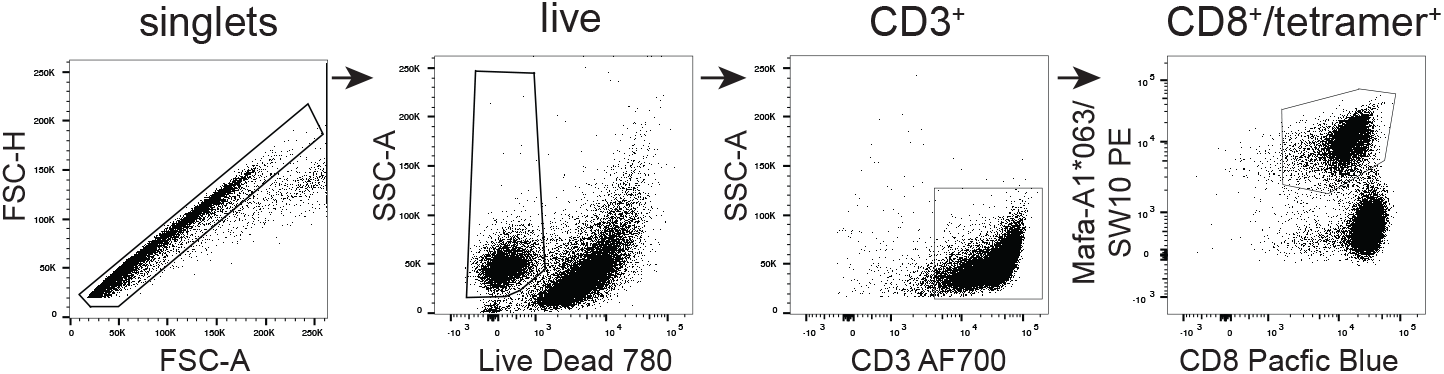
Tetrameric complexes of Mafa-A1*063/SW10 stain APOBEC3C SW10-specific CD8+ T cells. An APOBEC3C mHAg-specific T cell line was stained with PE-conjugated tetrameric complexes of Mafa-A1*063/SW10. Displayed is the flow cytometric gating strategy for data analysis, progressively selecting signlets, live cells, CD3^+^, and CD8^+^/tetramer^+^ cells. singlets, live cells, CD3^+^, and CD8^+^/tetramer^+^ cells.

### Tissue distribution of the APOBEC3C mHAg epitope

The tissue distribution of mHAgs can be exploited for therapeutic gain (30). This strategy is epitomized by the infusion of allogeneic T cells to treat hematologic cancers, which operates via the selective targeting of mHAgs expressed by recipient leukocytes, including malignant cells, to eliminate tumors without causing severe GvHD. We therefore sought to determine whether immune recognition of the SW10 epitope is similarly limited to the hematopoietic system, given that CD4^+^ T cells are known to express high levels of APOBEC3C.

To test nonhematopoietic cells, we collected skin explants and isolated fibroblasts from the APOBEC3C SW10^pos^ MCM, cy0743. In IFN-γ ELISpot assays, SW10-specific T cell lines isolated from the alloimmunized MCMs cy0738 and cy0747 displayed greater reactivity against the mHAg^pos^ BLCL from cy0743 compared with either the corresponding autologous BLCLs or mHAg^pos^ fibroblasts from cy0743, suggesting limited expression of SW10 outside the hematopoietic system (Figure 7). The APOBEC3C SW10 epitope may therefore serve as a model hematopoietic tissue-restricted antigen in MCMs.

**Figure 7.**
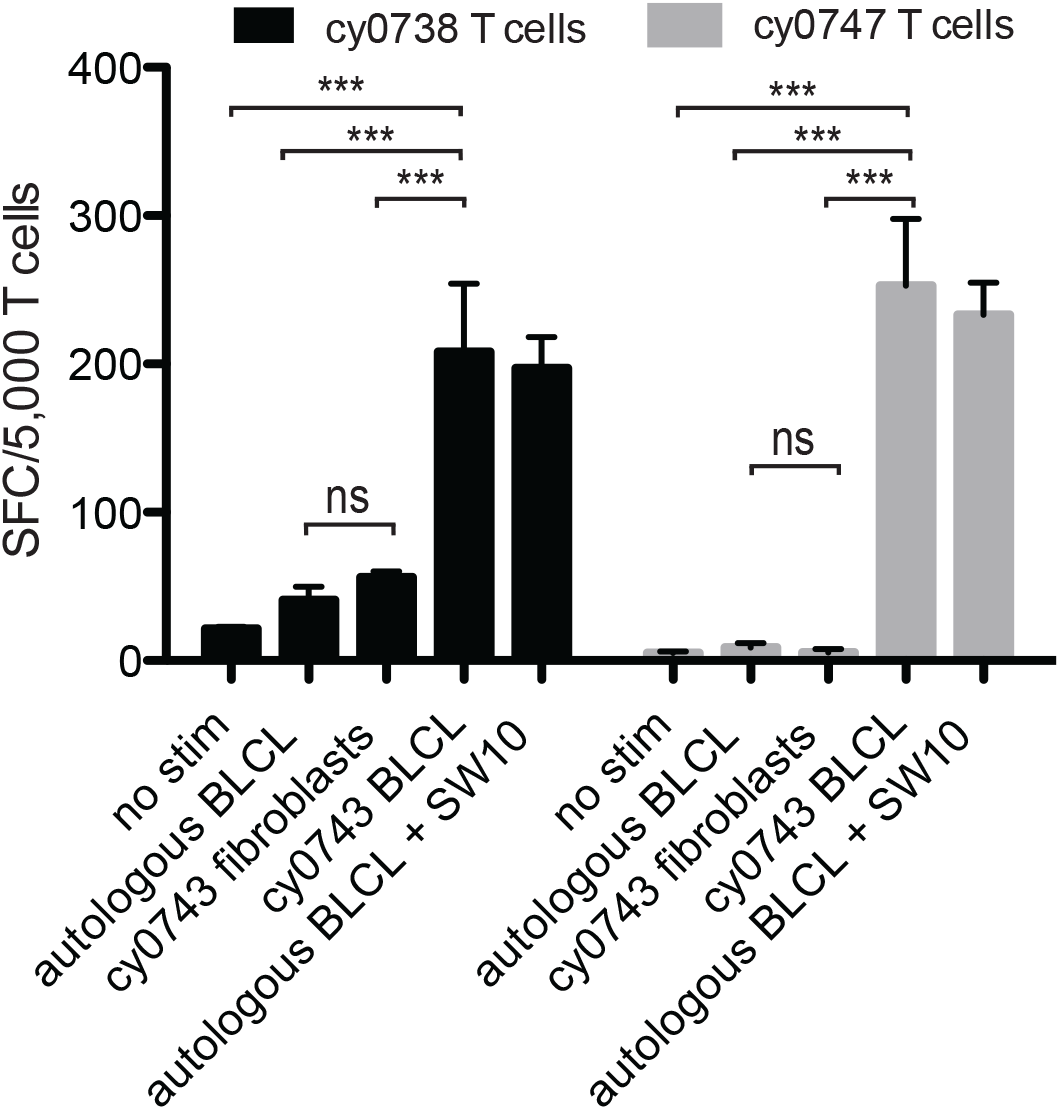
Tissue restriction of the mHAg epitope in APOBEC3C. APOBEC3C SW10-specific T cell lines from MCMs cy0738 and cy0747 were incubated with equal numbers of fibroblasts or BLCLs from the mHAg^pos^ animal cy0743 in an IFN-γ ELISpot assay. T cells cultured alone or incubated with equal numbers of the corresponding unpulsed autologous BLCLs were included as negative controls, and T cells incubated with equal numbers of the corresponding SW10 peptide-pulsed autologous BLCLs were included as a positive control. Data are shown as a mean ± SD for each condition. Differences between groups were assessed using a one-way ANOVA with Tukey's test for multiple comparisons. ***p<0.0001; ns=not significant. SFC, spot-forming cell.

### Frequency of the SW10-encoding SNP in 68 MCM

To determine the overall prevalence of the SW10-encoding SNP, irrespective of MHC haplotype, we expanded our analysis to include an additional 47 MCMs. Across the entire WES dataset, five animals had a SNP at position NC_027902.1:80990325 in *APOBEC3C*, representing a population frequency of 7.35% among 68 MCMs. Four animals had the G->T polymorphism at this position, encoding the leucine variant found in SW10. The other animal had a G->A nucleotide polymorphism at this position, encoding glutamine, which although not tested in this study, may similarly induce alloreactive T-cell responses in Mafa-A1*063-positive MCMs.

## Discussion

In this study, we used a combination of alloimmunization, WES, and bioinformatics to identify a mHAg in MCMs. Although a previous study described the conservation of human HA-1, HA-2, and H-Y mHAgs in chimpanzees and rhesus macaques (31), the epitope identity and immunogenicity of these mHAgs was not confirmed *in vivo.* Accordingly, the SW10 epitope in APOBEC represents the first clearly defined mHAg in NHPs.

Our investigation was made possible by the unique genetics of MCMs. In contrast to most outbred populations, MCMs passed through an artificial bottleneck approximately 400 years ago (32, 33), which resulted in a 23% reduction in the number of SNPs (33) and limited MHC diversity compared with mainland cynomolgus macaques (26, 27). These genetic peculiarities made it possible to assemble 21 MHC-identical animals, subdivide them into discrete groups based on mHAg expression, and map the SW10 epitope following the identification of SNPs in APOBEC3C.

Fortuitously, the MHC class I allomorph Mafa-A1*063 restricts APOBEC3C SW10. *Mafa-A1*063* is a “universal” A locus MHC-I allele expressed by the three most common MCM MHC haplotypes (M1, M2, and M3) (27). As a result, approximately 88% of MCMs have the potential to present APOBEC3C SW10. However, the SNP encoding this mHAg is present at a much lower frequency, estimated at only ~7% based on our analysis of 68 MCMs. Ideally, this SNP would be present at a higher frequency for ease of study. It should be noted that similar frequency limitations apply to MHC-homozygous MCMs (26). Although potentially limiting in the context of future interventional studies, breeding programs could be established to increase the frequency of APOBEC SW10 mHAg^pos^ MCMs.

*APOBEC3C* belongs to a family of well-known restriction factors that limit the replication of HIV and SIV. Apolipoprotein B mRNA editing enzyme catalytic polypeptide-like 3 (APOBEC) proteins are cytidine deaminases that protect against endogenous retroelements, retroviruses, and lentiviruses by inducing the hypermutation of viral genomes during reverse transcription (34). *APOBEC3* genes are expressed widely in immune cells (35–37). In line with these gene expression studies, we found that APOBEC3C SW10-specific T cells recognized BLCLs but not fibroblasts isolated from a mHAg^pos^ MCM. However, *APOBEC3C* is also expressed in a variety of human tissues, including the gut and the lungs (36, 37). The corresponding tissues may therefore be susceptible to attack by APOBEC3C SW10-specific T cells in MCMs. Our analysis nonetheless mirrors the screening process used to select tissue-restricted mHAg-specific T cell clones for infusion into patients with refractory leukemia (38), suggesting that the APOBEC3C SW10 epitope is sufficiently tissue-restricted to minimize the risk of severe GvHD in MCMs.

T cell immunotherapy is a useful and often curative strategy for treating hematologic cancers (38–47). A similar approach could potentially be used to target latent reservoirs of HIV (48, 49). Such reservoirs are limited to immune cells and are thought to be reduced or even eliminated by GvH responses after allo-HSCT (10–12). However, it is difficult to test this hypothesis in humans, because very few HIV+ patients undergo allo-HSCT. Moreover, different conditioning regimens are typically used in these rare cases, and efficacy likely depends to some extent on the receipt of HSCs with homozygous expression of CCR5Δ32. Our results may therefore facilitate the development of standardized models to investigate the antiviral effects of GvH responses in NHPs.

Our immunization strategy was adapted from human clinical trials investigating paternal lymphocyte immunotherapy (LIT) as a treatment for recurrent miscarriages (50, 51). In these studies, LIT was well tolerated and did not increase the risk of autoimmunity or GvHD (52, 53). In MHC-matched individuals, this alloimmunization strategy likely enriches for T cells targeting mHAgs expressed by leukocytes and could feasibly be extended to map mHAgs in other species, like outbred macaques or humans. Such *in vivo* enrichment protocols may offer advantages over traditional *in vitro* approaches for the identification of immunodominant mHAgs (54–57).

Further refinements to the methods described here could expedite the identification of mHAgs. A key limitation in our study was the inefficient generation of BLCLs. Although we generated WES data from 21 MHC-identical MCMs, we were only able to immortalize B cells from 10 animals, which constrained our ability to sieve candidate SNPs. The inclusion of a statistical component in the WES segregation analysis would also have likely enhanced the identification and prioritization of candidate SNPs (56). In addition, we focused our analysis on coding regions, which preclude the capture of intron-located or splice variant-generated SNPs (58). Whole-genome sequencing and/or mass spectrometric approaches would circumvent this particular limitation and allow the identification of such “cryptic” mHAgs.

In conclusion, we have identified a mHAg epitope in APOBEC3C restricted by the common MCM MHC-I allomorph Mafa-A1*063. Importantly, allorecognition of this epitope appeared to be limited to cells of hematopoietic origin, suggesting potential utility as a model antigen for testing mHAg-targeted immunotherapies in NHPs. In particular, we anticipate our discovery will allow the systematic analysis of GvH responses in the context of therapeutic interventions designed to combat latent immunodefficiency viruses, which are readily modeled in MCMs.

## Acknowledgments

We thank the veterinary staff at the Wisconsin National Primate Research Center (WNPRC) for their assistance. This study was funded by the National Institutes of Health (NIH) via grants R01 AI118495 and R24 OD021322 awarded to M.R.R. Additional support was provided by the Office of Research Infrastructure Programs/ OD via grant P51OD011106 awarded to the WNPRC at the University of Wisconsin-Madison. This research was conducted in part at a facility constructed with support from the Research Facilities Improvement Program via grants RR15459-01 and RR020141-01. D.A.P was supported by a Wellcome Trust Senior Investigator Award (100326/Z/12/Z). The funders had no role in study design, collection, analysis, interpretation of the data, or manuscript preparation.The content is solely the responsibility of the authors and does not necessarily represent the official views of the NIH.

## Author contributions

**Jason T. Weinfurter** performed experiments, acquired and analyzed data, and wrote the manuscript. **Michael E. Graham and Adam J. Ericsen** performed whole exome segregation analysis. **Lea M. Matschke** acquired and analyzed data. **Sian Llewellyn-Lacey and David A. Price** generated the Mafa-A1*063/SW10 tetramer. **Roger W. Wiseman** contributed to sample preparation and study methodology. **Matthew R. Reynolds** acquired funding, conceived and directed the project, and wrote the manuscript. All authors edited the manuscript and approved the final version.

## Materials and Methods

### Ethics statement and animal care

Cynomolgus macaques (*Macaca fascicularis*) used in this study were cared for by the staff at the Wisconsin National Primate Research Center according to the regulations and guidelines of the University of Wisconsin Institutional Animal Care and Use Committee, which approved this study (protocol g00695) in accordance with the recommendations of the Weatherall Report and the principles described in the National Research Council’s Guide for the Care and Use of Laboratory Animals. Macaques were housed in enclosures with at least 4.3, 6.0, or 8.0 sq. ft. of floor space, measuring 30, 32, or 36 inches high and containing a tubular polyvinyl-chloride or stainless-steel perch. Each enclosure was equipped with a horizontal or vertical sliding door, an automatic water lixit, and a stainless-steel feed hopper. Macaques were fed twice daily using a nutritional plan based on recommendations published by the National Research Council. Feeding strategies were individually tailored to the age and physical condition of each animal. Carbohydrate, energy, fat, fiber (10%), mineral, protein, and vitamin requirements were provided in an extruded dry diet (2050 Teklad Global 20% Protein Primate Diet). Dry diets were supplemented with fruits, vegetables, and other edible objects (e.g., nuts, cereals, seed mixtures, yogurt, peanut butter, popcorn, and marshmallows) to provide variety to the diet and to inspire foraging and other species-specific behaviors. To further promote psychological well-being, macaques were provided with food enrichment, human-to-monkey interaction, structural enrichment, and manipulanda. Environmental enrichment objects were selected to minimize the chances of pathogen transmission from one animal to another and from animals to care staff. Macaques were evaluated by trained animal care staff at least twice daily for signs of pain, distress, and illness by observing appetite, stool quality, activity level, and physical condition. Animals presenting abnormally for any of these clinical parameters were provided with appropriate care by attending veterinarians. Macaques were sedated with ketamine before each experimental procedure, and reversed with atipamezole after each experimental procedure. Animals were monitored regularly until fully recovered from anesthesia.

### Immunization of cynomolgus macaques

PBMCs were separated from EDTA-anticoagulated blood ia density gradient centrifugation using Ficoll-Paque PLUS (GE Healthcare). Freshly isolated PBMCs from four donor M1/M2 MCM were pooled and resuspended in RPMI-1640 containing 10% fetal calf serum (FCS; R10) and 5 ug/ml phytohemagglutinin-P (Sigma-Aldrich). The culture was incubated overnight at 37°C in a 5% CO2 atmosphere. Activated PBMCs were harvested the following morning, washed twice with phosphate-buffered saline (PBS), resuspended in PBS, and loaded into tuberculin syringes. Each macaque was then immunized with up to 4×10^7^ activated PBMCs as described previously (59).

### Growing bulk T cell lines to potential mHAgs

mHAg-specific T cell lines were generated from cryopreserved PBMCs isolated from alloimmunized macaques and thawed as described previously (60). Briefly, 5×10^6^ PBMCs were combined with 5×10^6^ irradiated BLCLs from MHC-identical donor MCMs in 5 ml of RPMI-1640 containing 15% FCS (R15) and 10 ng/ml recombinant human IL-7 (R&D Systems). Cultures were supplemented 2 days later with 1.25 ml of R15 containing 100 U/ml recombinant human IL-2 (R&D systems; R15-100). R15-100 was added every 2 to 3 days thereafter, and T cell lines were stimulated weekly with equal numbers of irradiated BLCLs from MHC-identical donor MCMs. Peptide-specific T cell lines were grown similarly using autologous irradiated BLCLs pulsed with 15-mers A3C-1 and A3C-B (GenScript).

### Limiting dilution cloning

Bulk T cell lines were counted with Trypan Blue exclusion dye and suspended at 1 cell/ml in R15-100. Each line was distributed across three 96-well plates at a mean of 0.2 cells/well and placed at 37°C in a 5% CO_2_ incubator. T cell clones were fed twice weekly by replacing half of the media volume with R15-100 and restimulated periodically with a mixture of irradiated MHC-identical donor BLCLs.

### ELISpot assays

ELISpot assays were performed using precoated monkey IFN-γ ELISpot^PLUS^ Kits according to the manufacturer’s instructions (Mabtech). Briefly, 5,000 clonal T cells and 5,000 fibroblasts or BLCLs derived from individual MHC-identical animals were added to each well in a total of 200 ul of R10. Positive control wells contained concanavalin A (Sigma-Aldrich). Alternatively, 5,000 bulk T cells and 5,000 fibroblasts or BLCs derived from individual MHC-identical animals were added to each well in 200 μl of R10. Positive control wells contained autologous BLCLs and the APOBEC3C SW10 peptide. All tests were performed in triplicate. Plates were incubated overnight at 37°C in a 5% CO_2_ atmosphere and imaged using an AID ELISpot Reader (Autoimmun Diagnostika GmbH). Differences in the number of IFN-γ spot-forming cells (SFCs) after stimulation with allogeneic versus autologous fibroblasts or BLCLs were assessed for significance using a one-way ANOVA with Tukey’s test for multiple comparisons in Prism version 5.0 (GraphPad Software Inc.).

### Mapping the APOBEC3C SW10 epitope

The mHAg epitope in APOBEC3C was mapped using a T cell line generated from an alloimmunized MCM in the presence of autologous irradiated BLCLs pulsed with the 15-mer peptides A3C-A and A3C-B (Genescript). The minimal optimal epitope was determined using overlapping 8-mer, 9-mer, 10-mer, and 11-mer peptides (Genescript). Briefly, 100,000 mHAg^neg^ BLCLs were incubated for 1 hour with each peptide at a final concentration of 1 uM and washed twice with R10. Next, 100,000 cells from the APOBEC3C T cell line were added to each tube in R10 containing 10 ug/ml brefeldin A (Sigma-Aldrich). Cells were then incubated for 5 hours at 37°C in a 5% CO2 atmosphere, washed twice with PBS, and stained with LIVE/DEAD Fixable Near-IR dye according to the manufacturer’s instructions (Thermo Fisher Scientific). To identify peptide-specific responses, cells were washed with R10, stained for 30 minutes at room temperature with anti-CD3 Alexa 700 (clone SP34-2; BD Biosciences) and anti-CD8 Pacific Blue (clone RPA-T8; BD Biosciences), washed twice with PBS containing 2% FCS (FACS buffer), fixed in 1% paraformaldehyde (PFA), permeabilized in FACS buffer containing 0.1% saponin, and stained for 1 hour at room temperature with IFN-γ FITC (clone 4S.B3; BD Biosciences). Cells were then washed twice with FACS buffer and resuspended in 1% PFA. Targeted peptides were assayed similarly across a range of concentrations to identify the most immunogenic epitope. Data were acquired using an LSRII flow cytometer (BD Biosciences) and analyzed using FlowJo software version 10 (FlowJo LLC).

### Segregation analysis

Segregation analysis was performed using a custom-built OSX application entitled Variant Segregator. This tool provides a user interface for setting up comparison groups, which allows the interrogation of multiple sample groups without the need to generate filtered vcf files using command line tools. The Variant Segregator identifies candidate SNPs using a simple scoring system that compares each animal in the mHAg^pos^ and mHAg^neg^ groups to the Mmul-8.0.1 rhesus macaque reference genome. The scoring system assigns 5 points for each macaque with a homozygous SNP mismatch relative to the reference genome, 3 points for heterozygous SNP mismatches, and 0 points for homozygous SNP matches. The final segregation score is the point sum in the mHAg^pos^ group minus the point sum in the mHAg^neg^ group. The segregation scores were used to identify and prioritize SNPs present in mHAg^pos^ MCMs for further analysis.

